# Fast and accurate estimation of species-specific diversification rates using data augmentation

**DOI:** 10.1101/2020.11.03.365155

**Authors:** Odile Maliet, Hélène Morlon

## Abstract

Diversification rates vary across species as a response to various factors, including environmental conditions and species-specific features. Phylogenetic models that allow accounting for and quantifying this heterogeneity in diversification rates have proven particularly useful for understanding clades diversification. Recently, we introduced the cladogenetic diversification rate shift model (ClaDS), which allows inferring subtle rate variations across lineages. Here we present a new inference technique for this model that considerably reduces computation time through the use of data augmentation and provide an implementation of this method in Julia. In addition to drastically reducing computation time, this new inference approach provides a posterior distribution of the augmented data, that is the tree with extinct and unsampled lineages as well as associated diversification rates. In particular, this allows extracting the distribution through time of both the mean rate and the number of lineages. We assess the statistical performances of our approach using simulations and illustrate its application on the entire bird radiation.

## 2 Introduction

Diversification rates vary widely across species, according to a variety of factors including species-specific features (e.g. life-history, phenotypic, or behavioral traits) or characteristics of the environment experienced by the species (e.g. resource availability, habitat heterogeneity). Understanding how and why these rates vary is central to understanding what makes species richness vary across species groups and geographic regions. Diversification rate heterogeneity is manifest in empirical phylogenies, which imbalance is typically much higher than expected under a homogeneous birth-death model (Aldous, 2001; Blum and François, 2006). Over the years, several phylogenetic approaches have been developed to estimate diversification rates from dated phylogenies of extant species and to relate these diversification rates to biotic and abiotic factors (Stadler, 2013; Pennell and Harmon, 2013; Morlon, 2014).

Recently, important efforts have been made to develop phylogenetic models of diversification that account for diversification rate heterogeneity (Alfaro et al., 2009; Rabosky, 2014; Barido-Sottani et al., 2018; Höhna et al., 2019; Maliet et al., 2019; Laudanno et al., 2020). Such models are required to avoid spurious inference of diversification histories linked to model misspecification (i.e. assuming homogeneous rates when this hypothesis is clearly violated, Rabosky, 2010; Morlon et al., 2011). They have also been developed to directly test the effect of various traits on diversification rates (see all the models of the SSE family, *e.g*. Maddison et al., 2007; FitzJohn, 2012), to identify major shifts in the tempo of clade diversification (Alfaro et al., 2009; Rabosky, 2014; Barido-Sottani et al., 2018), or to infer more subtle rate variations (Maliet et al., 2019).

In parallel to these developments, there has been an increasing interest in estimating species-specific diversification rates. Jetz et al. (2012) demonstrated the utility of such measures for understanding present-day geographic variation in species richness. In the later paper the authors introduced the DR measure, which can be computed for each extant species. DR has since been widely used to understand how diversification rates vary across gradients, such as elevational (Quintero and Jetz, 2018) and latitudinal gradients (Rabosky et al., 2018). It has also been used in tip-rate correlation tests, in order to assess the influence of traits on diversification rates, thereby providing an alternative to SSE models that can suffer from high false discovery rates (Rabosky and Goldberg, 2015; Harvey and Rabosky, 2017). Lineage-specific diversification rates have also been computed with BAMM (Bayesian Analysis of Macroevolutionary Mixtures, Rabosky, 2014), or similar models that have subsequently been developed to account for potential shifts in diversification rates in now-extinct lineages (Barido-Sottani et al., 2018; Höhna et al., 2019; Laudanno et al., 2020).

We recently developed the cladogenetic diversification rate shift model (ClaDS), which also provides species-specific estimates of speciation rates (Maliet et al., 2019). In contrast to other model-based approches (Rabosky, 2014; Barido-Sottani et al., 2018; Höhna et al., 2019; Laudanno et al., 2020) that capture rare shifts with large effects, ClaDS is specifically designed to capture more frequent shifts with small effects. A recent study showed that this ‘many small shifts’ model shows higher marginal likelihoods than the ‘few large shifts’ models when applied to bird clades (Ronquist et al., 2020). Rather than assuming that speciation rates are taken from a (usually small) set of rate regimes (Rabosky, 2014; Barido-Sottani et al., 2018; Höhna et al., 2019; Laudanno et al., 2020), ClaDS assumes that each lineage has its own speciation rate. At each speciation event, the two daughter lineages are assigned new speciation rates sampled in a distribution parameterized on the parental rate. This feature of ClaDS makes it particularly relevant in the context of estimating species-specific rates, as rates are explicitly modeled under an assumption of species-specificity rather than as coming from a limited set of rate regimes. It is also particularly interesting in the context of tip-rate correlation tests, as the power of such tests is limited by the number of distinct rate regimes present in a given phylogeny (Rabosky and Huang, 2016). Summary statistics such as DR also capture fine rate variations, however their exact relationship to diversification rates is unclear. DR for example is the inverse of the equal-split (ES) statistic initially developed in conservation biology to account for species evolutionary relatedness, which apportions the total phylogenetic branch-lengths of a tree among its constituent species (Redding and Mooers, 2006). Although DR has been widely used as a measure of diversification rates (Jetz et al., 2012; Rabosky et al., 2018; Title and Rabosky, 2019), the only mathematical link between this statistic and diversification rates that we are aware of is that DR converges to the speciation rate under a time-constant and homogenous model in the absence of extinction (i.e. the Yule model). Simulations under various models with heterogeneous rates have shown that ClaDS provides more accurate estimates of speciation rates than estimates from BAMM or DR (Maliet et al., 2019).

In our previous implementation for fitting ClaDS to empirical phylogenies, we sampled the parameters of the model and the species-specific diversification rates using a Metropolis Hastings Monte Carlo Markov Chain (MCMC) Bayesian framework. This requires multiple computations of the likelihood associated to a given reconstructed phylogeny (i.e. a phylogeny that includes a sample of only present-day species). In order to account for all the ‘hidden’ rate shifts happening at speciation events that are unobserved on the reconstructed phylogeny (i.e. all the speciation events that did not give rise to sampled descendants at present) in the computation of the likelihood, we need to numerically resolve a set of Ordinary Differential Equations (ODEs), which is computationally intensive. This computational cost limits the empirical applicability of ClaDS.

Here, we develop a much more efficient approach for fitting ClaDS to empirical phylogenies, using data augmentation (DA) within a MCMC in a Bayesian framework. DA is a classical statistic inference method that relies on introducing latent variables that simplify the evaluation of the model likelihood (Tanner and Wong, 1987). In the field of phylogenetic methods, DA has been used to map DNA substitution histories on phylogenies (Mateiu and Rannala, 2006; Lartillot and Poujol, 2011), to infer ancestral biogeographical histories (Landis et al., 2013), to assess the effect of a discrete character on trait evolution (May and Moore, 2020), or to develop biogeographic models with competition (Quintero and Landis, 2020). It has also been applied to estimate diversification rates, by augmenting the reconstructed phylogeny with extinct and unsampled lineages in a constant rate homogeneous birth-death model, in order to account for uniform (Bokma, 2008) and dispersed (Cusimano et al., 2012) species sampling.

We enhance the tree with extinct and unsampled lineages together with their lineage-specific speciation rates. This dramatically reduces the computational cost of ClaDS, because simulation is fast and the likelihood of the enhanced tree can be quickly computed analytically. In addition to making ClaDS much more computationally efficient, the DA implementation outputs distributions of the enhanced data; this provides estimates than can otherwise be hard to obtain, such as estimates of diversity through time (Morlon et al., 2011; Etienne et al., 2012; Manceau et al., 2019; Billaud et al., 2020), or of speciation rates through time. We test our new implementation of ClaDS using simulations and illustrate its application by fitting ClaDS to the entire bird radiation.

## 3 Methods

### 3.1 Model and notations

We develop our DA inference approach of ClaDS for the model referred to as ClaDS2 in Maliet et al. (2019), which is a model with constant turnover rate (i.e. extinction rate divided by speciation rate). We focused on this particular version of ClaDS as it is the one that produces trees with the most realistic tree shapes (Maliet et al., 2019), and best fit to empirical data (Ronquist et al., 2020). The model (illustrated in Figure 1 of Maliet et al., 2019) is based on the classical birth-death model (Nee et al., 1994), but each lineage *i* has its own speciation rate *λ_i_*. The process starts with a single lineage with speciation rate λ_0_. When lineage *i* speciates to give birth to two daughter lineages *s*_*i*,1_ and *s*_*i*,2_, the rates of the two new lineages are independently drawn in a lognormal distribution with parameters *log*(*αλ_i_*) and *σ*, where *α* is a deterministic trend parameter and *σ* controls the stochasticity of rate inheritance. We define *m* = *α* * *e*^σ^2^/2^ the expected relative change in speciation rate (this is the mean of the lognormal distribution). With *m* < 1, speciation rates tend to decrease at speciation, while with *m* > 1 they tend to increase. Lineage *i* can also go extinct at rate *ελ_i_*, where *ε* is the turnover parameter, assumed constant through time and homogeneous across lineages. Finally, lineages that survive to the present time are sampled independently from each other with probability *f*, with 0 < *f* <= 1.

**Figure 1:**
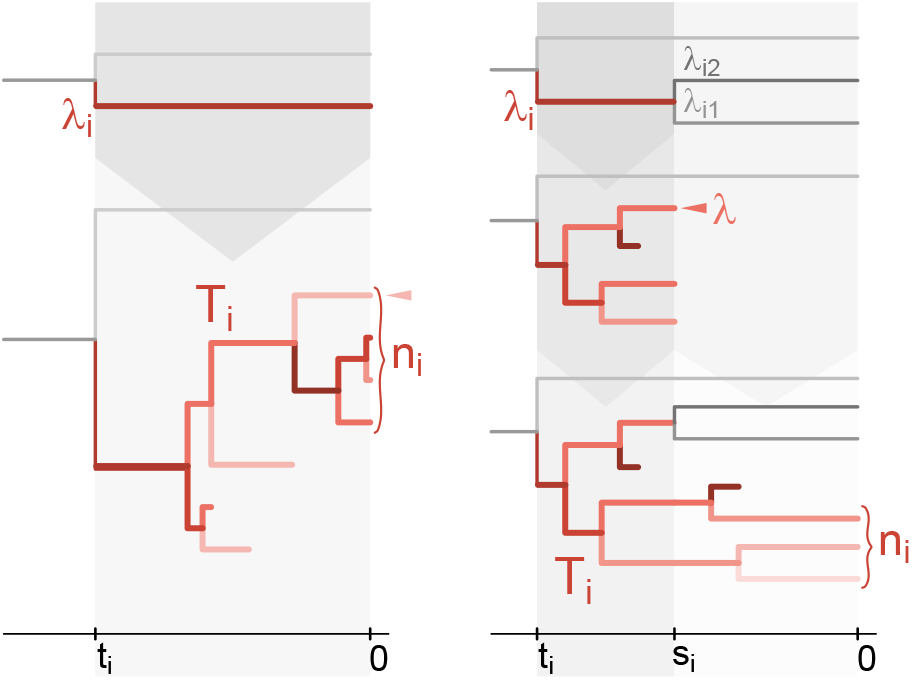
Illustration of the augmentation of a branch of the reconstructed phylogeny with extinct and unsampled lineages. Left: A terminal branch *i* starting at time *t_i_* with speciation rate *λ_i_* is augmented by simulating a phylogeny *T_i_* from the ClaDS model. If *T_i_* goes extinct before the present it is rejected; otherwise it is accepted with a probability proportional to the probability that exactly one lineage from *T_i_* is sampled at present, *n_i_f* (1 — *f*)*^n_i_^*. One tip of the accepted tree is sampled at random (pink arrow) to compute the tip rate of branch *i*. Right: An internal branch *i* starting at time *t_i_* with speciation rate *λ_i_* and branching at time *s_i_* with daughter rates *λ*_*i*1_ and *λ*_*i*2_ is augmented by first simulating a phylogeny from the ClaDS model with initial speciation rate *λ_i_* for a time *t_i_* — *s_i_*. If the process goes extinct before time *s_i_*, the proposal is rejected. Otherwise we sample at random one of the tips at *s_i_* (pink arrow, this tip has speciation rate *λ*), that will be the one branching in the reconstructed tree. For the other lineages alive at *s_i_*, we continue the simulation for a time *s_i_*. The resulting phylogeny *T_i_* is accepted with a probability proportional to the probability density of the two new rates times the probability that no lineage from *T_i_* will be sampled at present, *λν*(*λ, λ*_*i*1_)*ν*(*λ, λ*_*i*2_)(1 — *f*)^*n_i_*^.

### 3.2 Data dugmentation

We use data augmentation (DA) to improve the computational efficiency of the estimation of ClaDS parameters from dated reconstructed phylogenies, while integrating over all the possible rate changes along phylogenetic branches. DA allows sampling from the posterior distribution *p*(Θ|*X*) of the model’s parameters Θ knowing the observed data *X*, by introducing latent variables *Z* (also called the augmented data). Recursively simulating the latent variables Z knowing Θ (from the distribution *p*(*Z*|Θ,*X*)), then sampling the model’s parameters Θ knowing *Z* (from the distribution *p*(Θ|Z, *X*)) builds a Monte Carlo Markov Chain (MCMC) with stationary distribution *p*(Θ|*X*) (when marginalizing for Θ). By marginalizing the chain for the augmented data *Z*, we also have access to the distribution of the augmented data knowing the observed data, *p*(*Z*|*X*).

In our case, the observed data are any given reconstructed phylogeny; the model parameters are the four hyperparameters (*λ*_0_, *α, σ* and *ε*) and the speciation rates at the beginning of each branch; the augmented data are the unobserved speciation events and associated new speciation rates, as well as the extinction events. Augmenting the tree enables an efficient sampling of the parameters knowing the complete tree, because we can easily compute the likelihood of a complete tree under ClaDS analytically (see Appendix).

For the augmentation step, we simulate extinct and unsampled lineages for one branch from the reconstructed phylogeny at a time. This is accurate because lineages diversify independently from each other under the ClaDS model. Branches are updated using a Metropolis Hasting (MH) algorithm. In the MH algorithm, a new state *x* is proposed in replacement for the former state *x*\* by sampling from a proposal density *p*(*x*|*x*\*). The new state is accepted with probability 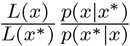, where *L*(.) is the posterior density. 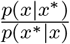 is called the Hastings ratio, and corrects for an eventual asymmetry in the proposal density *p*(*x*|*x*\*) ⪅ *p*(*x*\*|*x*).

At each augmentation step and for each branch, we replace the branch by a phylogeny with associated branch-specific rates (Fig 1), which is obtained by simulating a phylogeny under our model with the parameters from the MCMC at the step. This phylogeny might have only one edge if no event occurs during the simulation before the present (in the case of augmenting a terminal branch), or before the following branching event observed in the reconstructed tree (in the case of augmenting an internal branch). For a terminal branch of length *t_i_* with initial rate *λ_i_*, we simulate a phylogeny with initial rate *λ_i_* for a time *t_i_*. If the process goes extinct, the proposal is rejected. Otherwise if *n_i_* lineages survise to the present, it is accepted with probability min 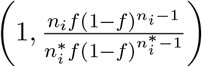 where 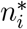 is the number of species that survived to the present in the augmented tree at the previous step for this branch (see Appendix). If the newly proposed augmented branch is rejected, we keep the previous one until the next augmentation step. Similarly, for an internal branch starting at time *t_i_* with speciation rate *λ_i_* and branching at time *s_i_*, we first simulate the process with a speciation rate *λ_i_* for a time *t_i_* — *s_i_*. If it goes extinct, the proposal is rejected. Otherwise, we select at random one of the lineages alive (that has speciation rate *λ*) to be the lineage that branches at time *s_i_*, and continue the process to the present for all the other alive lineages, which results in *n* >= 0 lineages alive at present time. The new augmented branch is accepted with probability min 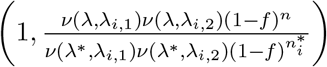 where 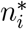 is the number of species that survived to the present for this branch in the previous augmented tree, and *λ*\* is the speciation rate for branch *i* at time *s_i_* in the previous augmented tree (see Appendix).

The probability density of each of the model parameters knowing the complete tree and all the other parameters can be readily obtained from the likelihood of the complete tree by fixing all the other parameters to their current value (see Appendix), so we use Gibbs sampling to update them. We sample each parameter from its density conditioned on the other parameter values using slice sampling (Neal, 2003).

### 3.3 Implementation

We implemented our functions in the PANDA package (https://github.com/hmorlon/PANDA. jl) in Julia (Bezanson et al., 2017). Our new implementation allows to specify different sampling probabilities for subclades in the phylogeny. In practice a user attributes a sampling probability to each tip (the sampling fraction of the subclade this species belongs to); our code attributes a sampling probability to each internal branch of the tree, taken to be the weighted mean of the subclades it subtends. More specifically, a branch with daughter clades that have *n_l_* and *n_r_* tips and sampling probabilities *f_r_* and *f_l_* is attributed the sampling probability 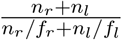.

To monitor the convergence of the MCMC, we run 3 chains simultaneously. Every *n_iter_* iterations, we compute the Gelman statistic (Gelman et al., 2014) for the 4 hyperparameters and the lineage-specific rates for each branch in the reconstructed phylogeny. We stop the algorithm when the Gelman statistics are below 1.05 for all the parameters.

As prior distributions, we use an inverse gamma prior with parameters (0.05, 0.5) for *σ*^2^ (we used an inverse gamma prior for *σ* in Maliet et al. (2019), but using it on *σ*^2^ is more natural, as the inverse gamma is the conjugate prior of the normal distribution with variance *σ*^2^). For ln(α) we use a normal distribution with parameters (—0.05,0.1), for *ε* a flat prior distribution on all positive values, and for the various speciation rates, including *λ*_0_, a flat prior distribution on the log scale.

In addition to the model parameters (hyperparameters and branch-specific rates for each branch from the reconstructed tree), we keep track of a number of summary statistics computed from the enhanced tree. We don’t record the enhanced trees at each MCMC step, which would be too heavy in terms of memory, but the function can output a sample from the complete tree distribution. We slice the time between the root of the phylogeny and the present into *n_timeslice_* equal intervals (which can be specified by the user and which we set at 50 as the default and for this study), and record, for each of the corresponding time points, the number of lineages and mean speciation rate over all the lineages in the enhanced tree. We also record the speciation rate at present for each tip in the reconstructed phylogeny. When sampling is incomplete (*f* < 1), there can be several tips in the augmented tree corresponding to a given extant species; in this case we pick one at random and record its rate.

The MCMC gives a sample from the posterior density of the model parameters, that can be summarized using a variety of metrics, including credibility intervals, means, and medians. We use the Maximum A Posteriori (MAP) for each of the parameters as point estimates. To do so we apply the function density from the R-package stats (R Core Team, 2013) to our MCMC chain, and record the parameter values for which the maximum density is reached. For the all the rates (*λ*_0_ and all the branch-specific rates), this is done after taking the logarithm of the chains, which we found to perform better. For the sake of memory use, only the initial rate of the branch is recorded at each MCMC iteration (for the terminal branch we also record the rate at present), so the inferred rates correspond to the rates at the beginning of the branch. If there is an interest in studying the rate variation along a given branch, this information can be retrieved from the sample of complete phylogenies outputted by the function. Point estimate for the number of lineage through time is obtained by taking the mean of the chain for each time point, as we found this estimate to perform better than the MAP for this quantity.

### 3.4 Test on simulated trees

We test the performance of our new inference method using simulations. We implement a new simulation algorithm of ClaDS conditioned on the number of tips; this new implementation avoids the bias linked to stopping the simulation as soon as the desired number of tips is reached (Hartmann et al., 2010) (see Appendix). In all simulations, we fix *λ*_0_ to 0.1; varying this parameter is equivalent to rescaling the branches of the trees but does not have an impact on tree shape. We draw the hyperparameters (*σ*, *α*) from their prior distributions, with the condition that the mean of the new rates *m* = *αe*^*σ*^2^/2^ < 1.3 to insure reasonable rate values and tree sizes. The prior distribution of ε is improper (i.e. it does not integrate to a finite value), so we draw *ε* from a uniform distribution on [0,1]. We test the model performances for sampling probabilities *f* in {1,0.9,0.5} and tree size *n* in {50,100, 200, 500}. For a given tree size *n* and sampling probability *f*, we start by simulating a complete tree conditioned on having ⌊*n*/*f*⌋ tips, where ⌊.⌋ is the floor function. We then keep each tip with equal probability *f*, conditional on the fact that at least one species from each of the two subtrees starting at the crown is sampled. To avoid numerical issues, we reject any tree with branches shorter than 1*e* — 10 or trees for which branch lengths span more than 6 orders of magnitude. These are very unrealistic trees and are unlikely to be encountered in empirical data. We simulate 500 trees for each combination of *n* and *f*, leading to a total number of 6000 simulated trees.

We run the inferences for our 6000 simulated trees, thinning every 30 iteration. We compute the Gelman statistics every 200 recorded iterations, after discarding the first quarter of the iterations. We stop the run as soon as the Gelman statistics are below 1.05 for all parameters (the hyperparameters and the branch-specific rates). In rare occasions one of the chain gets stuck in parameter space and the Gelman statistics does not go down; in this case we stop the run after 500.000 iterations even if the Gelman statistic is above the threshold value. This occurred in only 16 of our 6000 simulations, that we discarded from our analysis.

To assess the quality of the estimation of the hyperparameters, we compute MAP estimates and compare these estimates to the simulated values. To assess the quality of the estimation of the branch-specific speciation rates, we compute for each tree three different statistics: the correlation between simulated and estimated rates, the regression slope of the estimated versus simulated rates, and the mean relative error (exp(mean(log(*λ_infered_/λ_simultion_*)))). We compute these three statistics for the internal branches and for the tips.

Finally, we investigate the quality of inferred temporal dynamics of mean speciation rates and species richness by recording for each tree the number of time points (within the 50 time slices) for which the true values (directly computed from the simulations) fell outside the 95% credibility intervals of the inferred values. The mean rates at a given time are computed across all lineages alive at this time, including those that do not leave any sampled species.

### 3.5 Empirical application

We illustrate the utility of our new implementation by applying it to the bird phylogeny computed with molecular data from (Jetz et al., 2012) with the Hackett backbone, containing 6670 species. We use TreeAnnotator from the software Beast with the Common Ancestor option for node height (Bouckaert et al., 2019) to obtain a Maximum Clade Credibility (MCC) tree computed from a sample of 1000 trees from the posterior distribution. We fix the sampling fractions for each of the subtrees of the tree from Jetz et al. (2012) as the ratio between the number of species in the molecular phylogeny over that in the phylogeny including all bird species.

## 4 Results

### 4.1 Performance of ClaDS with a DA implementation

The new fitting procedure of ClaDS with data augmentation runs much faster than our previous approach (Maliet et al., 2019), a few hours instead of a few months for phylogenies with about 200 tips. This allowed us to test the performance of ClaDS under a much wider range of parameters, in particular with incomplete sampling and in the presence of extinction.

The hyperparameters are overall well inferred (Fig 2). The stochasticity parameter *σ* is slightly overestimated for low values and underestimated for high values but gets closer to the simulated value as the size of the phylogeny increases (Fig 2A, D). The trend parameter *α* is slightly biased toward 1 but also gets closer to the true value with increasing tree size (Fig 2B). The estimates of the turnover rate *ε* are highly variable but unbiased (Fig 2C). The hyperparameters are equally well inferred for all studied sampling probabilities (Fig S1). The coverage probabilities (the percentage of 95% credible interval that contains the parameter value used in the simulations) are slightly below 95% but approach this value for *α* and *ε* (Fig2D). For *σ* it is about 85%, and the simulated *σ* value is above the coverage interval approximately 15% of the cases. This might come from the fact that our simulated values are not exactly sampled from our prior (the coverage probability should be 95% only if the simulated values are sampled from the prior), because we reject trees with m > 1.3, which generally corresponds to high *σ* values. The coverage probability somewhat surprisingly does not improve with tree size (Fig 2D), but the size of the coverage intervals gets smaller as tree size increases (FigS2), reflecting a higher precision in our estimates.

**Figure 2:**
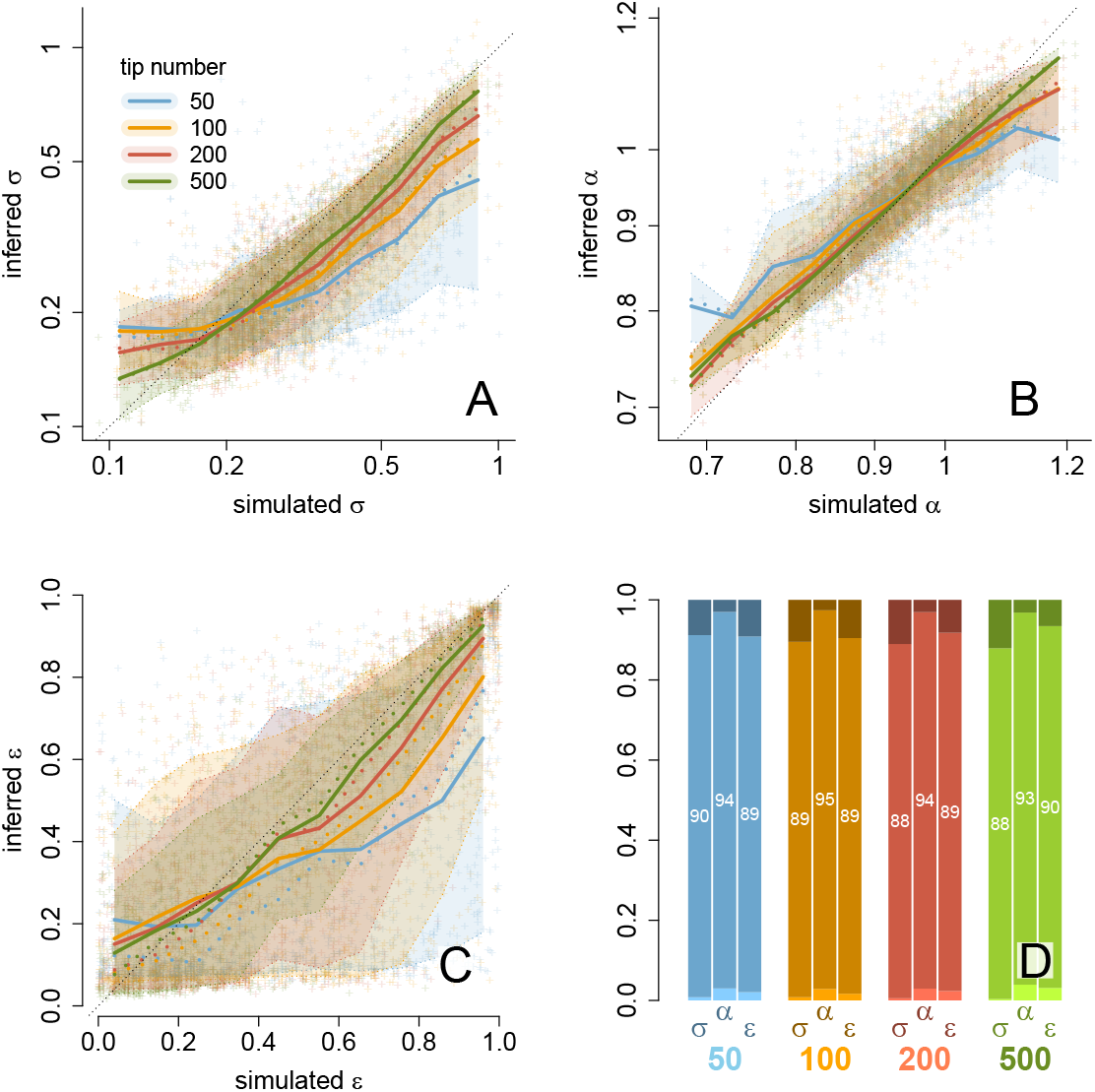
Estimation of ClaDS hyperparameters. A: Inferred versus simulated *σ* values on a log-log scale. Each cross is the result of an individual fit, and colors correspond to the tree sizes indicated in the legend. The thick solid lines display the sliding means, the thick dotted lines the sliding medians, and colored areas sliding intervals that contain 95% of the inferred values. The dotted black line shows the 1:1 bisector. B: Inferred versus simulated α values on a log-log scale. C: Inferred versus simulated *ε* values. D: Coverage probability for *σ*, *α* and *ε*. The middle of the barplot indicates the percentage of the 95% credibility interval that contains the simulated parameter value, while the darker and lighter bar respectively indicate the percentage of the inference for which the simulated value is lower or higher than the 95% credibility interval.

ClaDS returns branch-specific estimates for all branches of the augmented tree, as well as mean rates through time and diversity-through-time curves (Fig 3). The branch-specific speciation rates are also well inferred (Fig 4), as we see from the correlation between the simulated and inferred rates (Fig4A) and the mean relative error across simulated trees (Fig4B). The rates are slightly biased towards low values for small trees (Fig4, 50 tips) but this bias disappears quickly with increasing tree size, and the mean relative error is well centered on 1 for trees with 100, 200 and 500 tips. The correlation also improves with tree size (Fig 4A). For a given reconstructed tree size, the correlation is better when the clade is well sampled (Fig4A). This is also the case for the regression slope, that is generally below 1, because estimated rate variations tend to be smoothed with respect to the true variations (FigS3). It is the reverse for the mean relative error, which increases with the size of the complete tree (and thus is closer to 1 for lower *f*, Fig4B). This shows that the local variations are better inferred for well sampled trees, while the global trends are best inferred for big clades. The patterns are similar for tip rates (Fig4D,E).

**Figure 3:**
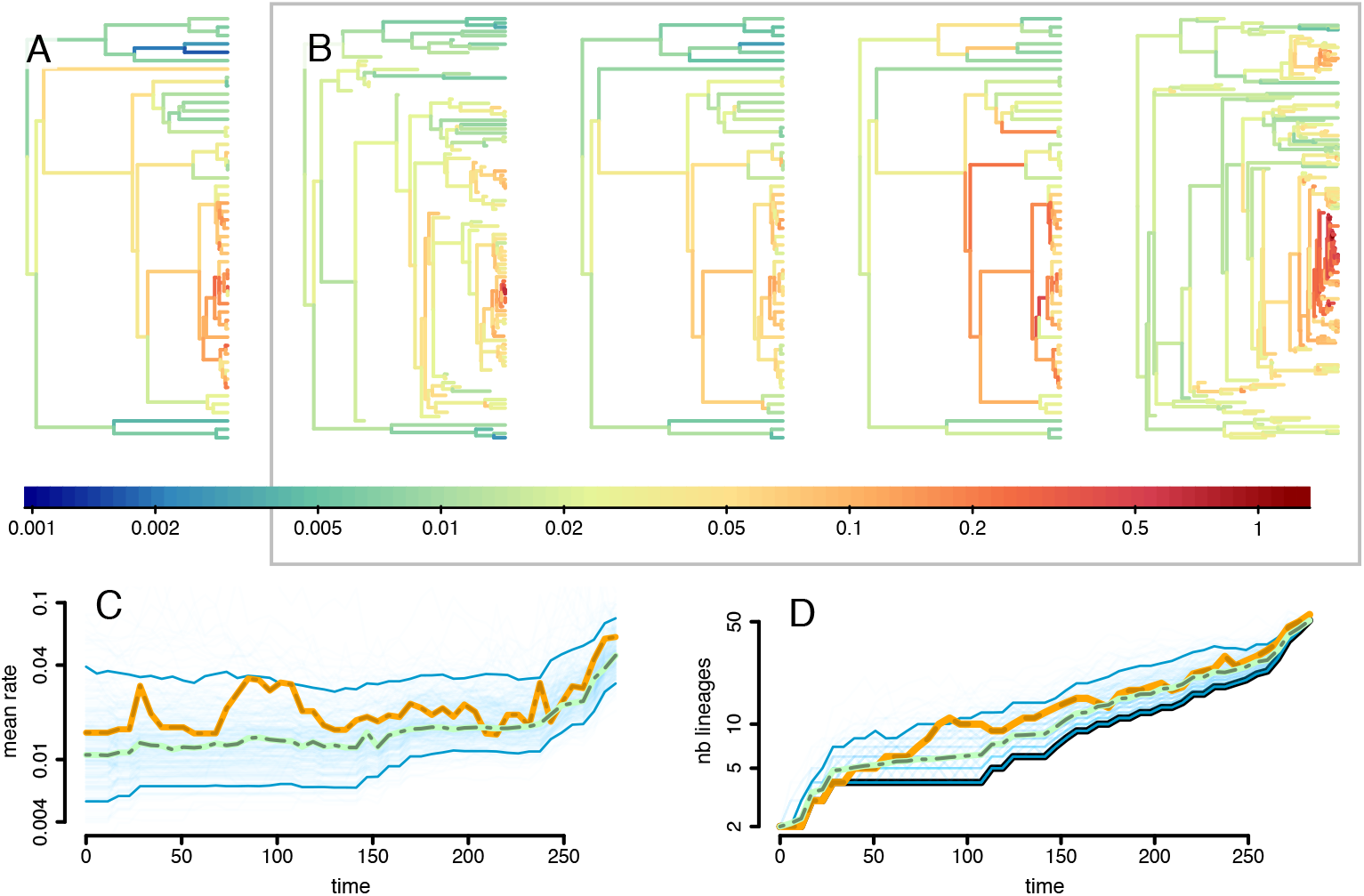
Inference on a simulated tree with 50 tips. A: The reconstructed simulated tree, with branches colored accorded to the simulated branch-specific speciation rates. B: Draws from the posterior distribution of complete phylogenies with speciation rates. C: Mean rate through time. Yellow line: true mean rate through time obtained in the simulation; Thin blue lines: individual draws from the MCMC; Thick blue lines: 95% credibility interval in the MCMC; Dotted green line: inferred mean rate through time estimated by taking the mean of the MCMC iterations. D: Diversity through time plot. Black line: number of lineages through time in the reconstructed phylogeny (the LTT plot); Yellow line: true number of lineages through time obtained in the simulation; Thin blue lines: individual draws from the MCMC; Thick blue lines: 95% credibility interval in the MCMC; Dotted green line: inferred number of lineages through time estimated by taking the mean of the MCMC iterations.

**Figure 4:**
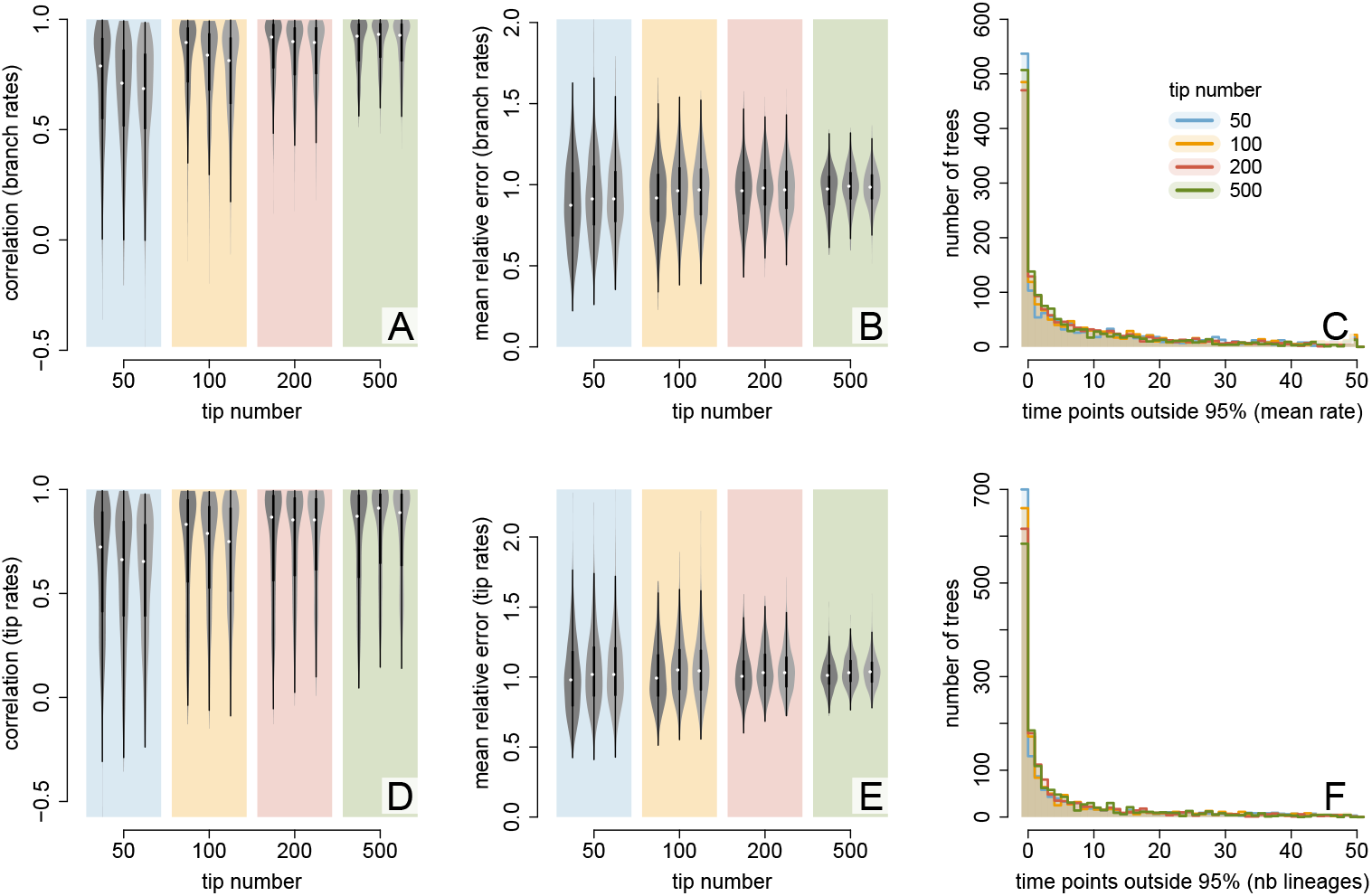
ClaDS performs well in estimating branch-specific rates, diversity-through-time, and mean rates through time. A: Correlation between simulated and inferred branch-specific speciation rates (the rates at the beginning of each branch) across all simulations, for each tip number and sampling probability (from left to right for each tree size: *f* = 1., 0.9, 0.5). The violin plots represent the distribution of correlation values. White dots represent the medians, and thick black lines the quartiles. B: Mean relative error in branch-specific speciation rates (the rates at the beginning of each branch). C: Distribution of the number of time points when the true curve of the mean speciation rate through time is out of the credible interval for each simulated tree. D: Correlation between simulated and inferred branch-specific speciation rates at present. E: Mean relative error in branch-specific speciation rates at present. F: Distribution of the number of time points when the true curve of the number of lineages through time is out of the credible interval for each simulated tree.

For about 30% of the trees, the simulated mean speciation rates fall within the inferred 95% interval over the entire time span of the simulation (Fig4C); for most of the trees for which this is not the case, they fall within the credibility interval at least a 80% of the times (Fig4C). For about 40% of the trees, the simulated number of lineages fall within the inferred 95% interval over the entire diversity-through-time curve (Fig4D); for most of the trees for which this is not the case, they fall within the credibility interval at least 80% of the times (Fig4D). Similarly to what happens for the hyperparameter, tree size increases the precision of the estimation but not coverage probability.

### 4.2 Illustration on the bird phylogeny

Our new implementation of ClaDS allows fitting the model with extinction and incomplete sampling to the entire bird phylogeny (6670 tips, Fig. 5), which we were not able to do in our previous study as it would have required a huge amount of time. We fit the full tree in less than a day with the new implementation, while in our previous study it took several months to approach convergence for family-level fits. Consistent with what we found on individual bird clades (Maliet et al., 2019), we find that the variability in rates is high, as indicated by the *σ* value (Fig. 5D, *σ* = 0.55), and branch-specific speciation rates are spread over more than two orders of magnitude (Fig. 5C, from 0.01 to 4 events per million years). We also confirm that there is a general tendency for rates to decline through time, as we can see from the trend parameter *α* (Fig. 5E, *α* = 0.68) and the mean new rate *m* = *α* * *e*^*σ*^2^/2^ (Fig. 5G, *m* = 0.79). Contrary to what could have been expected from these hyperparameter estimates, the estimated mean speciation rate appears relatively constant through time, with a slow decline for the first 60 million years, followed by a slow increase (Fig. 5A). We find a very low level of extinction (Fig. 5F, *ε* = 0.07). Consequently, the diversity-through-time curve is very similar to the LTT plot, diverging only close to the present because of incomplete sampling (Fig. 5B).

**Figure 5:**
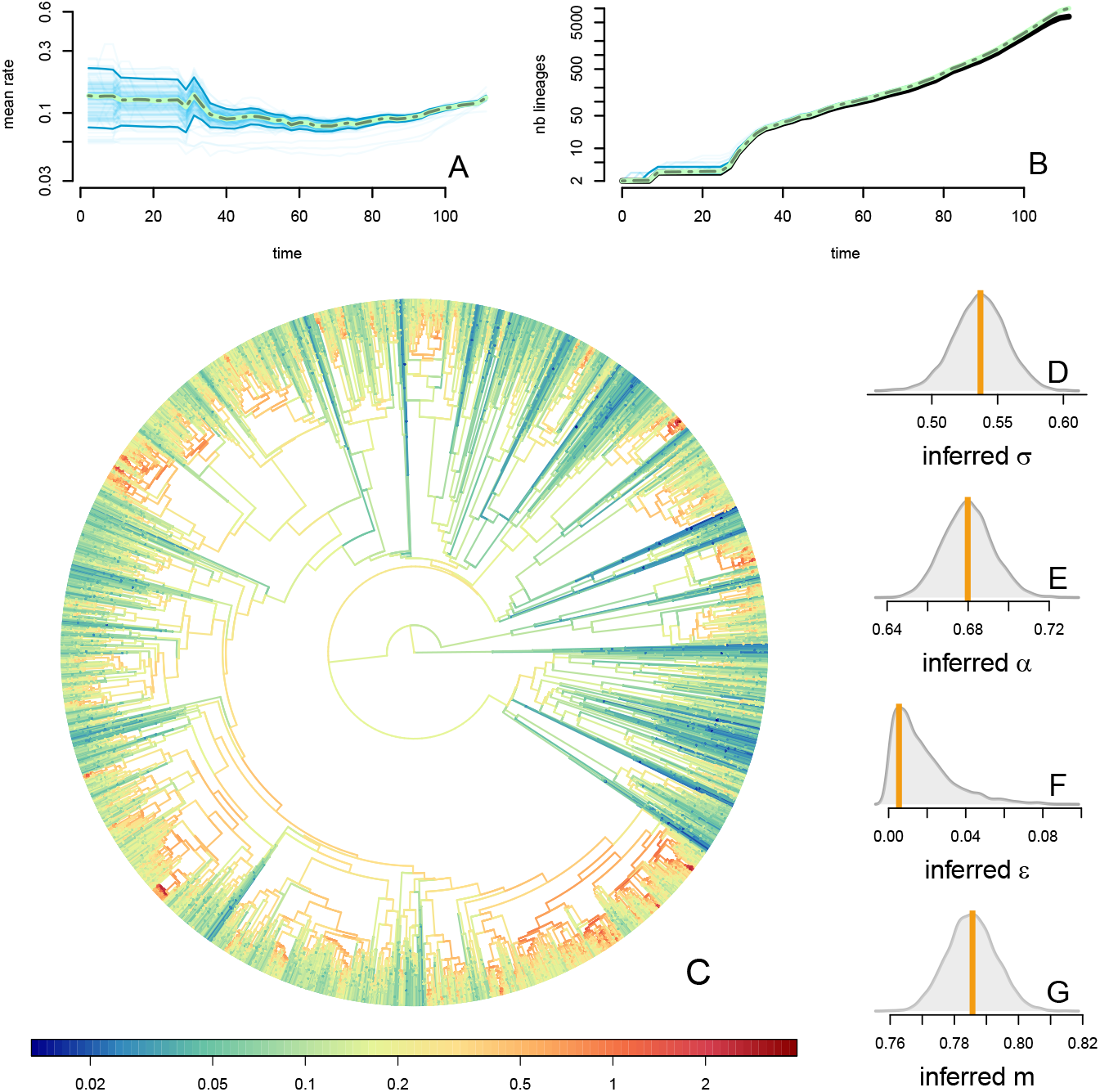
Fit of ClaDS to the entire bird radiation. A: Inferred mean rate through time, with individual MCMC iterations (thin blue line), the 95% credibility interval for each time point (thick blue line), the mean for each time point (dotted green). Time goes from root time to the present. B: Inferred number of lineages through time, with the count from the reconstructed phylogeny (black line), individual MCMC iterations (thin blue line), the 95% credibility interval for each time point (thick blue line), the mean for each time point (dotted green). Time goes from root time to the present. C: Inferred lineage-specific speciation rates for the bird phylogeny. D: Marginal posterior of the heterogeneity parameter *σ*. The orange line indicates the value of the MAP estimate. E: Marginal posterior of the trend parameter *α*. F: Marginal posterior of the turnover rate *ε*. D: Marginal posterior of the mean change in rate *m* = *αe*^*σ*^2^/2^.

## 5 Discussion

We developed a new implementation of ClaDS, using data augmentation, that allows a fast and accurate inference of branch-specific speciation rates, while accounting for extinction and incomplete sampling. This new implementation renders the application of ClaDS to large phylogenies feasible and should be useful for empirical studies that aim to understand how and why diversification rates vary across lineages. The approach provides more resolution than models with few large rate shifts (Alfaro et al., 2009; Rabosky, 2014; Barido-Sottani et al., 2018; Höhna et al., 2019; Laudanno et al., 2020) and more theoretical grounding than summary statistics such as DR (Jetz et al., 2012). We implemented our approach in a Julia package with a user-friendly interface, and we provide a tutorial illustrating how to run the model (SI).

In addition to estimating branch-specific speciation rates, our new implementation of ClaDS provides a distribution of complete phylogenies consistent with the empirical reconstructed tree, along with associated branch-specific rates (i.e. the augmented data). This can be useful to reconstruct past dynamics, as we have illustrated here with diversity-through-time and rate-through-time curves; we show that these curves are well estimated for most of our simulated trees. Other methods that allow the estimation of diversity-through-time curves from reconstructed phylogenies assume that diversification rates are homogeneous across lineages (Etienne et al., 2012; Manceau et al., 2019), or require an a priori specification of nodes in the phylogeny where rate shifts occurred (Morlon et al., 2011; Billaud et al., 2020). Almost all these methods also require estimating diversification rates (e.g. by maximum likelihood; but see Manceau et al., 2019) prior to computing the curves, while ours estimate diversification rates and diversity jointly along the MCMC. Our implementation thus allows estimating diversity-through-time curves while accounting for heterogeneous speciation rates among lineages and uncertainty in model parameters.

Since we compute mean rate-through-time curves from the complete, data-augmented phylogenies, we avoid potential biases due to not accounting for extinct or unsampled lineages by computing these curves on the reconstructed phylogeny (Rabosky, 2014). As illustrated here with the application on the bird phylogeny with an inferred *m* below 1 and a rather constant mean rate-through-time curve, the mean temporal tendency can be counter-intuitive given inferred *m* values. Indeed, even though *m* < 1 indicates the tendency for daughter rates to be lower than parental rates, lineages that by chance have a high rate have a higher probability of speciating and having offspring, such that the mean rate does not necessarily decrease through time. Hence examining mean rate-through-time curves in addition to *m* values is an essential step when studying the diversification dynamics of a clade with ClaDS.

Computing likelihoods for augmented data is in general much easier mathematically, and efficient computationally, than computing likelihoods for reconstructed data. Hence we can envision several extensions of the ClaDS model using data augmentation. For example, the current implementation can accommodate only bifurcating trees, but trees with polytomies could be accommodated in the future by simulating unresolved nodes during the tree enhancement phase, in the spirit of polytomy resolvers (Kuhn et al., 2011). Similarly, non-ultrametric trees, such as those that integrate extinct (fossil) species (Stadler, 2010), could readily be accommodated. It would also be rather straightforward to modify the distribution in which daughter rates are drawn to allow for both large and small rate shifts, which would allow testing how much variations in diversification rates are associated with major shifts, for example linked to key innovations, and small variations, for example linked to gradual trait evolution. Other potential extensions include integrating the effect of environmental correlates (Condamine et al., 2013; Lewitus and Morlon, 2017), species interactions (Etienne et al., 2012), or phenotypic evolution (Maddison et al., 2007) within ClaDS, which would allow testing the role of these various factors on diversification while accounting for rate heterogeneity.

Our approach shares some characteristics with the implementation of ClaDS by Ronquist et al. (2020) using Probabilistic Programming Languages (PPL) and alive particle filters. The two approaches rely on intensive simulation of the process. The use of particle filters has the additional advantage of giving an estimate of the marginal likelihood of the model, which is useful for model comparison. It would be interesting to compare the performance of both approaches in terms of computation time.

The fit of ClaDS to the full bird radiation, made possible by the new implementation, confirms our earlier results based on family-level analyses, and allows a comparison with previous studies that have used the same bird phylogeny (Jetz et al., 2012; Rabosky, 2016). The extent of rate variation we found with ClaDS is similar to what has been found with the DR statistic (Jetz et al., 2012) and wider than the one found when using BAMM (Rabosky, 2016). Interestingly, our rate through time curve is very similar to the one found for the same phylogeny in Jetz et al. (2012) (Fig 1), even though the model they used was very different from ours, with diversification rates that are homogeneous among lineages but can jump at given time points (Stadler, 2011). Consistently with previous studies (Jetz et al., 2012; Maliet et al., 2019; Ronquist et al., 2020), we infer a very low level of extinction in this tree. This could be due to the dating of this tree which was done using a pure birth prior, as dating priors can have a strong impact on tree dating and posterior diversification analyses (Condamine et al., 2015). It could also be due to the yet-to-be explained difficulty of inferring extinctions from empirical trees (Rabosky, 2010); as we have shown extinction is well inferred on trees simulated under the ClaDS process.

The fit of ClaDS to the full bird radiation reveals a clear tendency for decrease in rates at speciation (*m* = 0.78), as is regularly found in empirical phylogenetic studies (Phillimore and Price, 2008; Moen and Morlon, 2014). However, this does not translate into a rate decrease through time at the global level; we even find a slight increase in rates starting from 60 Myrs ago. The rate decline at speciation is compensated by the lineage selection effect (lineages with relatively high speciation rates leave more daughter lineages), which is all the more preponderant that *σ* (i.e. the level of stochasticity) is large. So even though a slow-down can be seen at a small phylogenetic scale, repeated bursts of diversification rates hide this signal at the global scale, as proposed by Henao Díaz et al. (2019). This result emphasizes the importance of using heterogeneous diversification models when studying large clades, which our new implementation now makes possible for ClaDS.

## 6 Appendix

### 6.1 Probability density of the complete phylogeny

The probability density of the complete tree with branch-specific rates is the combination of: the probability density of the waiting times between branching events ((1+*ε*)*λ*_*i*_*e*^-(1+*ε*)*λi*^), the probability that a given event is an extinction (*ε*/(1 + *ε*)) or a speciation (1/(1 + *ε*)), the probability density of the two new speciation rates after a branching event (*ν*(*λ_i_*, *λ*_*s*_*i*, 1__)ν(*λ_i_*, *λ*_*s*_i_, 2_)), the probability that no event happened on a given branch (*e*^-(1+*ε*)*λi*^). Calling *E_s_* the set of internal branches, it can thus be written as:

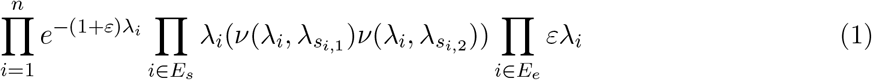

### 6.2 Updating a branch

We note Θ the model hyperparameters, *T_r_* the reconstructed tree, *T* the complete tree enhanced with *T_i_* at branch *i* of the reconstructed tree, *n_i_* the number of tips of *T_i_* alive at present time, and *T_i_* the complete tree without branch *i*. We also note *t_i_* the starting time of branch *i* in the reconstructed tree, and *s_i_* its end time. Finally, we note *λ_i,j_* (with *j* ∈ [1,*ns_i_*]) the branch-specific speciation rates of *T_i_*, with *λ_i,1_* = *λ_i_* the branch-specific speciation rate of branch *i* in *T_r_*. We use the same notations with a * upper script when referring to these quantities in the precedent MCMC state.

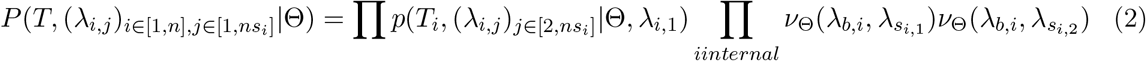

The probability to have *T_r_* as a reconstructed tree knowing the complete tree *T* is the probability that no species from *T_i_* was sampled if *i* is an internal branch ((1 — *f*)^*n_i_*^) and that exactly one species from *T_i_* was sampled if i is a terminal branch (*n_i_f*(1 — *f*)^n_i_-1^). So

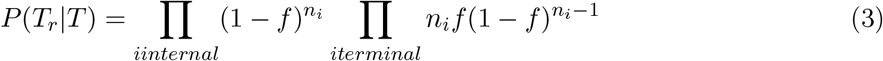

For branch *i*, we propose a new phylogeny *T_i_* simulated with ClaDS2 for a time *t_i_* with initial speciation rate *λ_i_*. So the probability to propose *T_i_* when the current enhanced tree is *T*\* is

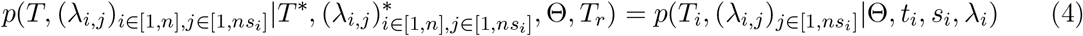

which is independent of the previous state 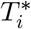, so the Hastings ratio is

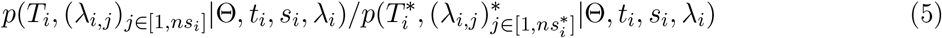

The posterior probability of the new state is

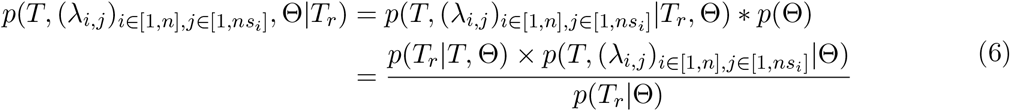

The new proposed branch is accepted with probability

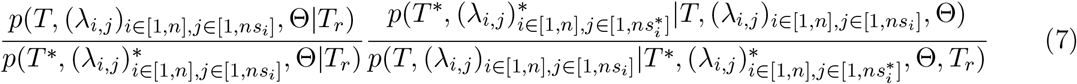

Injecting the respective value of these quantities, we obtain that the acceptance probability is, if branch *i* is internal

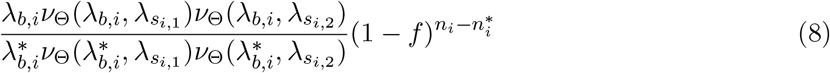

And, if branch *i* is terminal

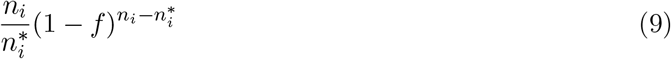

### 6.3 Simulating a phylogeny conditioned on tip number under ClaDS

In our former implementation of simulations of ClaDS conditioned on the number of tips at present (Maliet et al., 2019), we simulated a tree with *n* tips by running the model until *n* species were reached and stopping the simulation at the time of the following event. While this approach is very commonly used, it leads to biased tree sampling as soon as the model differs from the pure birth model, because it does not allow to simulate phylogenies in which the number of species has been higher than *n* before decreasing (Hartmann et al., 2010). Here we implement an alternative approach, the General Sampling Approach (GSA), that avoids this bias (Hartmann et al., 2010). The GSA consists in simulating the process until it is very unlikely that it will reach n species latter on, either because the process went extinct or because a larger number of species was reached (in our model, we typically stop when 5*n* species are reached). The simulation time is then drawn uniformly in the set of times during which there has been *n* species, and the corresponding tree is retrieved.

## Acknowledgments

The authors are very grateful to Leandro Arístide, Julien Clavel, Carmelo Fruciano, Sophia Lambert, Benoît Perez-Lamarque, Isaac Overcast, Ignacio Quintero, Ana Catarina Silva and Guilhem Sommeria-Klein for their helpful comments on an earlier version of this manuscript. This work was supported by the Labex MemoLife (postdoctoral funding to OM) and the European Research Council (grant ERC CoG-PANDA attributed to HM).

