## Supplementary Information for "Fast and accurate estimation of species-specific diversification rates using data augmentation"

November 2, 2020

Odile Maliet[a,*], Hélène Morlon[a]
[a] Institut de biologie de l'Ecole normale supérieure (IBENS), Ecole normale supérieure, CNRS, INSERM, PSL Research University, 75005 Paris, France

#### 1 Supplementary figures

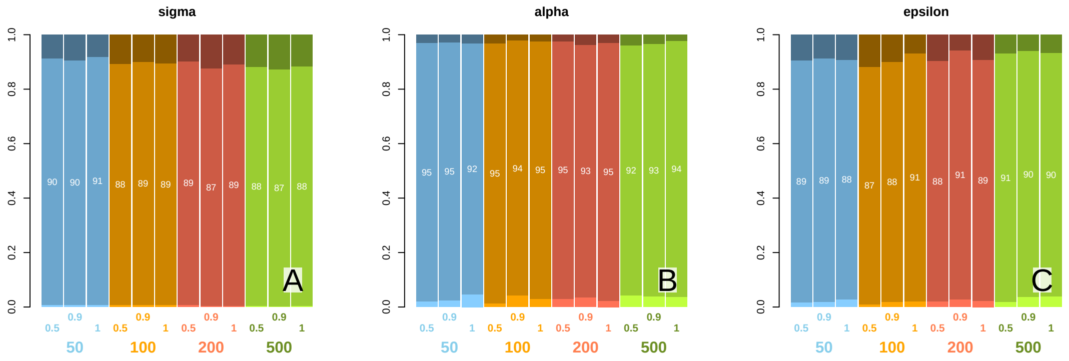


Figure S1: **Coverage probability of ClaDS hyperparameters.** A: Coverage probability for $\sigma$, for each tree size (50, 100, 222 and 500) and sampling probability (0.5, 0.9, 1.0). B: Coverage probability for $\alpha$, for each tree size and sampling probability. C: Coverage probability for $\varepsilon$, for each tree size and sampling probability. The middle of the barplot indicates the percentage of the 95% credibility interval that contains the simulated parameter value, while the darker and lighter bar respectively indicate the percentage of the inference for which the simulated value is lower or higher than the 95% credibility interval .

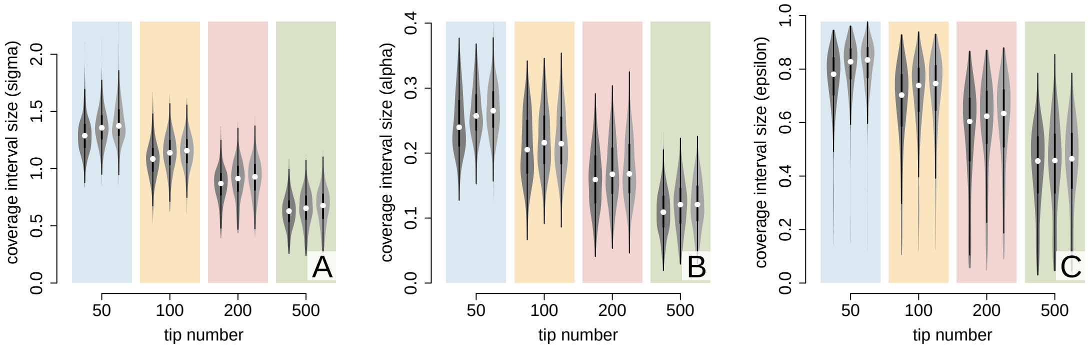


Figure S2: **Size of the coverage probability interval of ClaDS hyperparameters.** A: Size of the coverage probability interval for $\sigma$, for each tree size (50, 100, 222 and 500) and sampling probability (0.5, 0.9, 1.0). B: Size of the coverage probability interval for $\alpha$, for each tree size and sampling probability. C: Size of the coverage probability interval for $\varepsilon$, for each tree size and sampling probability. The violin plots represent the distribution of correlation values. White dots represent the medians, and thick black lines the quartiles. The different shades of gray of the violons display the different sampling probability (from left to right for each tree size: $f = 1., 0.9, 0.5$).

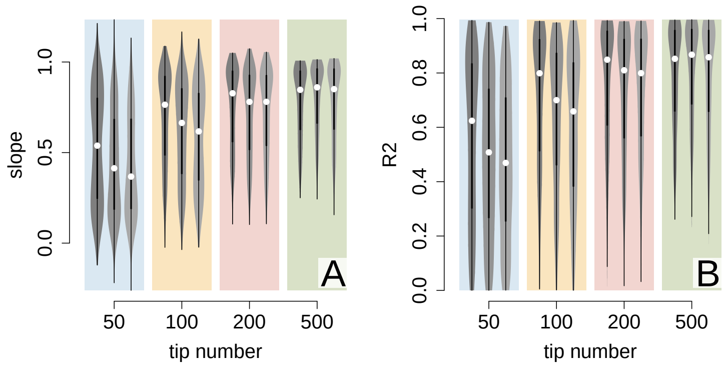


Figure S3: **Regression between simulated and inferred branch-specific speciation rates.** A: Distribution of the regression slopes, for each tree size (50, 100, 222 and 500) and sampling probability (0.5, 0.9, 1.0). B: Distribution of the $R^2$, for each tree size and sampling probability. The dots represent the medians, and thick black lines the quartiles. The different shades of gray of the violons display the different sampling probability (from left to right for each tree size: $f = 1., 0.9, 0.5$).

### 2 Tutorial

ClaDS is implemented in the `PANDA.jl` julia package (https://github.com/hmorlon/PANDA.jl). Here we show how to run the model on a tree and exploit the results. The most up-to-date version of this tutorial can be found on the package help page (https://hmorlon.github.io/PANDA.jl/stable).

#### 2.1 Installation

You need to download Julia following the instructions from
https://julialang.org/downloads/
The package can then be installed from a Julia session with the following command:

```
using Pkg
Pkg.add("PANDA")
```

PANDA uses R functions and packages for plotting. If you want to be able to use the plotting functions, the R language (R Core Team, 2013) needs to be installed on your computer. You will also need a few R packages to be installed, including : ape (Paradis et al., 2004), coda (Plummer et al., 2006), RColorBrewer (Neuwirth and Neuwirth, 2014), fields (Douglas Nychka et al., 2017). You can install them from a R session by typing

```
install.packages("ape", "coda", "RColorBrewer", "fields")
```

`PANDA` can then be loaded to a Julia session with

```
using PANDA
```

#### 2.2 Loading a tree

You can import a phylogeny to the environment using the `load_tree` function. Currently supported extensions include `.tre` and `.nex`.

```
my_tree = load_tree(tree_path)
```

#### 2.3 Running the inference

The parameter inference is ran with the function `infer_ClaDS`:

```
output = infer_ClaDS(my_tree)
```

You can save the result with

```julia
using JLD2
@save the_path_you_want_to_save_the_result output
```

and load it back to a julia session with

```julia
using JLD2
@load the_path_you_want_to_save_the_result output
```

**Incomplete sampling** By default, the function considers that the clade was perfectly sampled, i.e. that all the species alive at present time are included in the phylogeny. If it is not the case, the sampling fraction can be specified through the keyword argument `f`. `f` can be a `Float`, in which case the sampling fraction is taken as homogeneous on the whole phylogeny.

```julia
output = infer_ClaDS(my_tree, f = 0.94)
```

Alternatively, different sampling fractions can be specified for different subclades. To do so, `f` should be passed as a `loat64ArrayF` of length `n`, where `n` is the number of tip in the phylogeny. `f[i]` is the sampling fraction of the subclade that contains tip `i`. If the `Tree` object has tip labels (which can be accessed using `tip_labels(my_tree)`, the sampling fractions in `f` are in the same order as the tip labels, and `f[i]` is the sampling fraction of the subclade that contains the tip with label `tip_labels(my_tree)[i`'.

In the following example, the left subtree of `my_tree` is assigned the sampling fraction `0.3` and its right subtree the sampling fraction `0.8`.

```julia
#=
 create a vector of size n_tips(my_tree) where the first
 n_tips(my_tree.offsprings[1]) elements are equal to 0.3
 and the rest to 0.8
=#
f = [ i < n_tips(my_tree.offsprings[1]) ? 0.3 : 0.8 for i in 1:n_tips(my_tree)]

output = infer_ClaDS(my_tree, f = f)
```

#### 2.4 Result

The result is a `CladsOutput` object, that contains the following fields:

`tree`: the `Tree` object on which the inference was performed.

- `chains`: the resulting mcmc chains.
- `rtt_chains`: the mcmc chains of the mean rate through time.
- `σ_map`, `α_map`, `ε_map`, `λ0_map`: the MAP estimates of the model's parameters.
- `λi_map`, `λtip_map`: the MAP estimates of the branch-specific and present rates.

- `time_points`: Time points at which the number of lineages through time and rate through time are computed. The number of time points can be specified using the keyword argument `ltt_steps`.

- `DTT_mean`: Estimate of the number of lineages through time.

- `RTT_map`: Estimate of the mean rate through time.

- `enhanced_tree`: Sample from the complete phylogeny distribution. Their number can be specified through the keyword argument `n_trees`.

- `gelm`: Evaluation of the gelman statistics.

If the tree has tip labels, the tip rate for species `sp_name` can be extracted using the function `tip_rate`:

```
tip_rate(output, sp_name)
```

The result can be saved as a `.Rdata` object with the function `export_in_R`, so it can be manipulated in R.

**Plot the branch specific rates** It can be plotted using the `plot_CladsOutput` function. By default, this function plots the reconstructed phylogeny painted with the inferred branch-specific speciation rates, but other methods are available.

```
plot_CladsOutput(output)
```

Options for plotting the tree can be passed as a ‘String‘ to the ‘options‘ keyword argument. All the options of the ‘R‘ function ‘plot.phylo‘ from the ‘ape‘ package are available.

```
plot_CladsOutput(output, options = "type = 'fan'")
plot_CladsOutput(output, options = "lwd = 1, direction = 'leftwards'")
```

**Diversity through time plot** Using the keyword argument `method = "DTT"`, the function plots the estimate of the number of lineages through time.

```
plot_CladsOutput(output, method = "DTT")
```

On this plot, we have:

- black line: the LTT plot (number of lineages through time in the reconstructed phylogeny)

- thin blue lines: individual MCMC iterations

- thick blue lines: the 95% confidence interval

- dotted green line: the point estimates

**Mean rate through time plot** Using the keyword argument `method = "RTT"`, the function plots the estimate of the mean speciation rate through time.

```
plot_CladsOutput(output, method = "RTT")
```

Similarly to the diversity through time plot, we have here:

- thin blue lines: individual MCMC iterations
- thick blue lines: the 95% confidence interval
- dotted green line: the point estimates

**Marginal posterior densities** With the keyword argument `method = "density"`, the functions plot the marginal posterior density of a given model parameter or a summary statistics. Similarly, `method = "chain"` allows plotting the mcmc chains for this parameter.

```
plot_CladsOutput(output, method = "density")
plot_CladsOutput(output, method = "chain")
```

What parameter to plot is specified through the keyword `id_par`, for both `method = "density"` and `method = "chain"`.

```
plot_CladsOutput(output, method = "density", id_par = "sigma")
plot_CladsOutput(output, method = "density", id_par = "σ")

plot_CladsOutput(output, method = "density", id_par = "alpha")
plot_CladsOutput(output, method = "density", id_par = "epsilon")
plot_CladsOutput(output, method = "density", id_par = "lambda0")
```

The branch specific speciation rate of branch `i` is accessed with `id_par = "lambda_i"` or `id_par = "λ_i"`

```
plot_CladsOutput(output, method = "density", id_par = "lambda_5")
plot_CladsOutput(output, method = "density", id_par = "λ_2")
```

The tip rate of tip `i` is accessed with `id_par = "lambda_tip_i"` or `id_par = "λtip_i"`. Alternatively, if the tree has tip labels, the rate can be accessed using the species name with `id_par = lambda_tip_spname`.

```
plot_CladsOutput(output, method = "density", id_par = "lambdatip_2")
plot_CladsOutput(output, method = "density", id_par = "λtip_3")

sp_name = tip_labels(my_tree)[10]
id_par = "λtip_$(sp_name)"
plot_CladsOutput(output, method = "density", id_par = id_par)
```

Finaly, the rate through time and diversity through time can be accessed with `id_par = rate_time` and `id_par = div_time`, where `time` is an integer between `1` and `length(output.time_points)`.

```
plot_CladsOutput(output, method = "density", id_par = "rate_12")
plot_CladsOutput(output, method = "density", id_par = "div_33")
```
